# Phosphorylation of the ER-plasma membrane tether E-Syt2 facilitates the loss of contact sites in mitosis

**DOI:** 10.64898/2026.11.21.768271

**Authors:** Jawaher Al Hamaq, Kripa S Keyan, Fang Yu, Lubna Zarif, Khaled Machaca

## Abstract

Mitosis is associated with dramatic remodeling of intracellular organelles that supports their equal distribution between daughter cells. Recent investigations into membrane contact sites (MCS) show that they also remodel in mitosis with a significant loss of ER-plasma membrane CS (ERPMCS). However, the mechanisms governing this loss are unknown. Here we show that phosphorylation of the ERPMCS tether extended synaptotagmin 2 (E-Syt2) facilitates the loss of ERPMCS in mitosis. Preventing E-Syt2 phosphorylation in mitosis rescues the loss of ERPMCS. The persistence of ERPMCS is associated with delayed growth and progression through mitosis. This shows that E-Syt2 mitosis specific phosphorylation regulates its tethering potential and that the loss of ERPMCS is important for timely cell division.

## Introduction

Cell division is a fundamental process for multicellular organism growth, development, and homeostasis. Disruption of the tight regulation of mitosis can lead to disastrous consequences such as cancer, developmental defects, and tissue degeneration. In addition to the proper segregation of the genetic material, mitosis encompasses dramatic organelle remodeling to ensure their equal distribution to the two daughter cells to support future growth and divisions ^1^.

The partitioning of organelles to the two daughter cells occurs either in a directed fashion, stochastically, or a combination of both ^1–4^. Stochastic inheritance involves fragmentation of the organelle to allow for a balanced distribution into the two daughter cells. This is particularly important for single copy organelles such as the Golgi, but also occurs with larger organelle networks such as the mitochondria. Another central concept in organelle inheritance during mitosis is that most organelles cannot be generated de novo, implying that organelle inheritance encodes epigenetic information in their structure that regulates cell division ^2,3^.

The area of the plasma membrane (PM) is reduced during mitosis due to inhibition of constitutive exocytosis, which also affects the structure of the ER and Golgi ^2,3,5^. The ER adopts a more cortical position in mitosis away from the spindle, with the preponderance of the evidence supporting a more tubular structure ^1,6^. The nuclear envelope, which is continuous with the ER, breaks down resulting in the redistribution of inner envelope proteins to the bulk ER ^4,6–8^. This redistribution has been implicated in the remodeling of mitochondria-ER CS (MERCS) in mitosis. MERCS expand in mitosis leading to improved mitochondria respiration due to enhanced Ca^2+^ coupling between the ER and mitochondria ^9,10^. The inner nuclear envelope protein Lamin B receptor (LBR) enriches at MERCS in mitosis, which suggests it acts as a mitosis-specific MERCS tether ^10^.

The remodeling during mitosis affects not only the organelles themselves but also their interactions with each other. Inter-organelle communication and coordination is important for cellular homeostasis and uses either diffusible messengers, vesicular transport, or direct physical interactions through Membrane Contact Sites (MCS). MCS are close physical inter-organelle contacts where the membranes of two organelles are physically tethered <30-50 nm apart. This tight apposition favors rapid signaling between the two organelles, which has been best studied in the context of Ca^2+^ signaling and non-vesicular lipid transfer ^11–14^. MCS are stabilized by tethering proteins that bridge the two membranes. Tethering is typically through protein-protein interaction between tethers on the opposing membranes or through protein-lipid interactions and is best studied in the context of the ER as the largest organelle ^11,15^. In this paper we focus on ER-PM contact sites (ERPMCS), which are stabilized by various tethers, including VAPs, E-Syts, Junctophilins, Nir2/3, ORP5/8, GRAMD1a/2a, STEP, TMEM24, Kv2.1, and STIM1 ^11,15–18^.

Interest in the fate of ERPMCS in mitosis arose from investigating the mechanisms responsible for the inactivation of store-operated Ca^2+^ entry (SOCE) in mitosis. SOCE is a primary Ca^2+^ influx pathway activated downstream of various PLC-linked agonists and requires direct physical interaction at ERPMCS between the ER Ca^2+^ sensors STIM1/2 with the Ca^2+^ selective plasma membrane channels Orai1/2/3 ^19^. SOCE is invariably lost during cell division ^20,21^, yet the molecular mechanisms involved are not well defined. The phosphorylation of STIM1 during mitosis was implicated in SOCE downregulation, however its causality could not be validated ^22–24^. Yu et al. suggested that the primary mechanism responsible for the loss of SOCE in mitosis is the decrease in ERPMCS. They assessed ERPMCS density using ERPMCS markers and by direct quantification at the ultrastructural level ^25^. They further showed that the reduction in ERPMCS in mitosis was not associated with a global reduction in the expression of ERPMCS tethers.

Here, we focus on the role of one of the ERPMCS tethers, E-Syt2, in regulating ERPMCS in mitosis. E-Syt2 is an ER-anchored ERPMCS tether with glycerolipid transfer activity. It is one of three E-Syts with an SMP domain and multiple C-terminal C2 domains that bind to PI(4,5)P_2_ at the PM ^26,27^. E-Syt2 and 3 bind constitutively whereas E-Syt1 is Ca^2+^-dependent. E-Syts in addition transport diacylglycerol (DAG) from the PM to the ER ^28,29^. Double (E-Syt2/3) and triple E-Syt (E-Syt1/2/3) knockout mouse lines are viable and fertile, arguing for redundancy in E-Syts function at the whole organism level ^30^ ^31^.

## Results

### E-Syt2 mitosis-specific phosphorylation mutants

Because ERPMCS are lost in mitosis and because phosphorylation cascades play critical roles in mitosis entry and progression, we were interested in testing whether phosphorylation of ERPMCS tethers regulates their tethering capacity. Dephoure et al. performed a proteome wide screen for phosphorylated peptides in mitosis ^32^. We mined their results for the phosphorylation state of ERPMCS tethers, which revealed increased phosphorylation of several tethers (Table 1). We thus expressed several of those tethers to ascertain their distribution (Fig. 1A). Of the tested tethers, E-Syt2 and TMEM24 showed the most consistent ERPMCS enrichment (Fig. 1A). We chose to focus on E-Syt2 to determine whether its phosphorylation at mitosis-specific residues affects its ability to tether the ER to the PM. In mitosis E-Syt2 is phosphorylated at S/T 612, 623, 629, 667, 677 (Table 1, Fig. 1B). We thus generated two mutants with these S/T residues mutated to either Ala (E-Syt2-A) to generate a non-phosphorylatable E-Syt2 mutant; or to Glu (E-Syt2-E) to mimic phosphorylation at these sites with the negatively charged Glu side chain. All three clones had a N-terminal mCherry tag and when expressed localized to ERPMCS (Fig. 1C).

**Figure 1.**
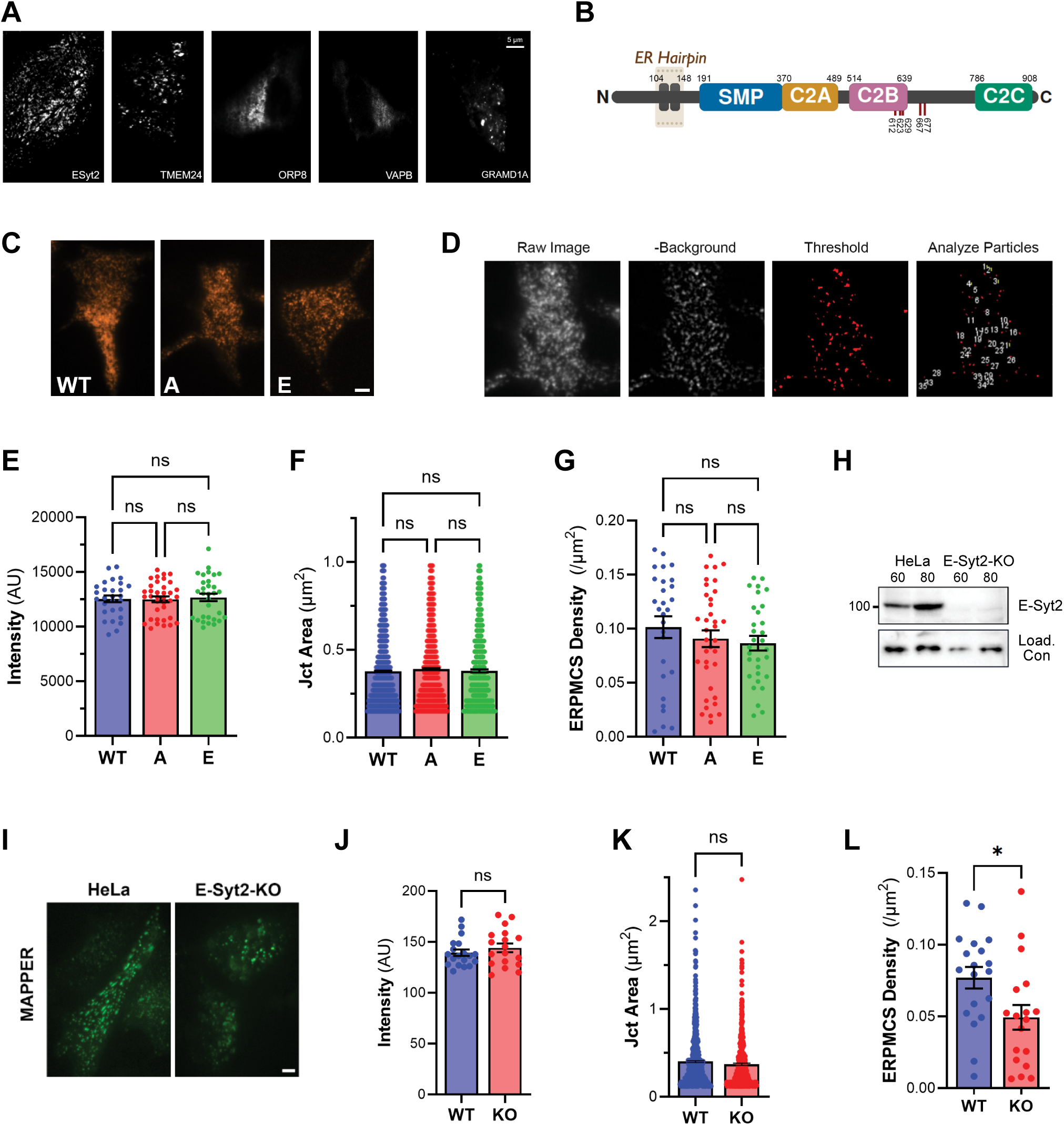
E-Syt2 phosphorylation mutants and knockout. (A) TIRF microscopy images of HeLa cells transiently expressing mCherry-E-Syt2, TMEM24-mCherry, mCherry-ORP8, GFP-VAPB and mCherry-GRAMD1A. Scale bar 5µm. (B) Schematic showing the E-Syt2 domain structure. Ser/Thr residues that are phosphorylated in mitosis are indicated by the red lines with the residue number. (C) Representative TIRF images of interphase HeLa cells transiently expressing E-Syt2-WT, non-phosphorylatable E-Syt2-A mutant and phosphomimetic E-Syt2-E mutant. Scale bar 5µm. (D) Example images from the ImageJ macro developed for automated quantification of ERPMCS. (E-G) Quantification of ERPMCS. (E) Intensity on a per cell basis, n = 28-36. (F) Individual junction area, n = 568-1894. (G) ERPMCS Density per cell, n = 28-36. Mean ± SEM; Anova with Tukey correction; ns not significant. (H) Western blot validation of E-Syt2 knockout HeLa cells. (I) Representative TIRF images of WT and E-Syt2 knockout HeLa cells expressing GFP-MAPPER. Scale bar 5µm. (J-L) Quantification of GFP-MAPPER-labeled ERPMCS. (J) Intensity per cell, n = 18-19. (K) Individual junction area, n = 529-823. (L) ERPMCS density per cell, n = 18-19. Mean ± SEM; unpaired t-test; * p < 0.05, ns not significant.

**Table 1.** Peptides from ERPMCS tethers with increased phosphorylations in Mitosis (Adapted from Dephoure et al.)

| Protein | G1 | M | Residue (Dephoure) | eSyt2 (NM_020728) | Peptide |
|---|---|---|---|---|---|
| ESYT1 |  | 1 | 959 |  | L**T**HVDSPLEAPAGPLGQVK |
|  |  | 3 | 963 |  | LTHVD**S**PLEAPAGPLGQVK |
| E-Syt2 |  | 1 | 660 | 612 | **S**HMSGSPGPGG**S**NTAPS**T**PVIGGSDKPGMEEK |
|  |  | 1 | 671 | 623 | **S**HMSGSPGPGG**S**NTAPS**T**PVIGGSDKPGMEEK |
|  |  | 1 | 677 | 629 | **S**HMSGSPGPGGS#NTAPS**T**PVIGGSDKPGMEEK |
|  |  | 4 | 715 | 667 | SSSSLLA**S**PGHISVK |
|  |  | 2 | 725 | 677 | SSSSLLASPGHISVKEP**T**P**S**IASDI**S**LPIATQELR |
| GRAMD1A |  | 1 | 364 |  | GHV**T**PNL**S**R |
|  |  | 1 | 368 |  | GHV**T**PNL**S**R |
|  |  | 1 | 636 |  | GLL**S**GLR |
| ORP3 |  | 1 | 16 |  | LV**SP**SRS**T**S**S**CSSK |
|  |  | 1 | 21 |  | LV**S**PSRS**T**S**S**CSSK |
|  |  | 1 | 23 |  | LVS#PSRS**T**S**S**CSSK |
|  |  | 4 | 372 |  | QLMEQDASS**S**PSAQVIGLK |
|  | 2 | 4 | 410 |  | IHAESLLLD**S**PAVAK |
| ORP8 |  | 2 | 63 |  | DLHQP**S**LSPA**S**PHSQGFER |
|  |  | 2 | 65 |  | DLHQPSL**S**PA**S**PHSQGFER |
|  |  | 4 | 68 |  | DLHQPSL**S**PA**S**PHSQGFER |
| TMEM24 |  | 2 | 411 |  | ALGPAATMAVELHYEEG**S**PR |
|  |  | 1 | 417 |  | NLG**T**PT**S**STPRPSITP**T**KK |
|  |  | 1 | 420 |  | NLG**T**PT**S**STPRPSITP**T**KK |
|  |  | 1 | 430 |  | NLG**T**PT**S**STPRPSITP**T**KK |
|  |  | 1 | 464 |  | LD**SP**SR**S**PSKVEVTEK |
|  |  | 1 | 468 |  | LD**S**PSR**S**PSKVEVTEK |
| VAPB |  | 1 | 146 |  | IISTTA**S**KTE**T**PIVSK |
|  |  | 9 | 150 |  | TE**T**PIVSK |
|  |  | 2 | 154 |  | TE**T**PIV**S**KSLSSSLDDTEVKK |
|  |  | 2 | 156 |  | **S**LSSSLDDTEVKK |
|  |  | 2 | 158 |  | SL**S**SSLDDTEVKK |
|  | 1 | 5 | 160 |  | SLSS**S**LDDTEVKK |

The reduction in ERPMCS in mitosis could be due to degradation of the tethers. We had previously tested several ERPMCS tethers in mitosis and did not detect any significant decrease but did not test E-Syt2. We therefore checked whether the expression of E-Syt2 is modulated in mitosis using SILAC (Stable Isotope Labeling by Amino Acids in Cell Culture) labeling ^25^. The levels of E-Syt2 were similar during both stages of the cell cycle, in contrast to STIM1, which was significantly downregulated in mitosis (Fig. S1A).

To quantify the tethering capacity of the E-Syt2 constructs, we assessed their localization to ERPMCS using a macro developed in image J to automate the process and minimize bias. In the raw images ERPMCS were typically not well defined with significant background as would be expected given the depth of the confocal slice (Fig. 1D), so we performed background subtraction followed by thresholding and particle analysis to quantify the area and density of ERPMCS (Fig. 1D). As the constructs were transiently transfected in HeLa cells, we used cells that express in the same range (Fig. 1E) to avoid any artifacts due to over expression; and then normalize on per cell expression level. We did not observe any difference in the junction area (Fig. 1F) or the density of ERPMCS on a per cell basis (Fig. 1G) between the three clones in HeLa cells transiently transfected.

We were concerned that endogenous E-Syt2 may mask changes in ERPMCS following the expression of the mutants, especially given the redundancy inherent in the three E-Syts and the multiple ERPMCS tethers. We therefore generated an E-Syt2 knockout HeLa cell line (E-Syt2-KO) to allow for expression of only the phosphomimetic (E-Syt2-E) or the non-phosphorylatable (E-Syt2-A) forms. We confirmed the loss of E-Syt2 using two different antibodies (Fig. 1H and S1B). Furthermore, we functionally tested the effect of the loss of E-Syt2 by quantifying the density of ERPMCS. Loss of E-Syt1/2 or all three E-Syts has been shown to decrease the density of ERPMCS ^27,33^. To determine whether E-Syt2 knockdown alone has the same effect, we used MAPPER as an ERPMCS marker ^34^ (Fig.1I). Knockout of E-Syt2 did not affect MAPPER intensity at the junctions (Fig. 1J) or ERPMCS area (Fig. 1K) but did significantly decrease ERPMCS density (Fig. 1L).

### Functional characterization of the E-Syt2 mutants in interphase

We then used colocalization of the different E-Syt2 constructs with MAPPER in the E-Syt2-KO cell line as a measure of their tethering capacity. Both the wild-type (WT) and the E-Syt2-A mutant colocalized with MAPPER with similarly high PCC (Pearson’s Correlation Coefficient), whereas the phosphomimetic mutant, E-Syt2-E, showed reduced co-localization, suggesting a diminished stability at ERPMCS (Fig. 2A and 2B). This would be consistent with the hypothesis that E-Syt2 phosphorylation decreases its tethering capacity.

**Figure 2.**
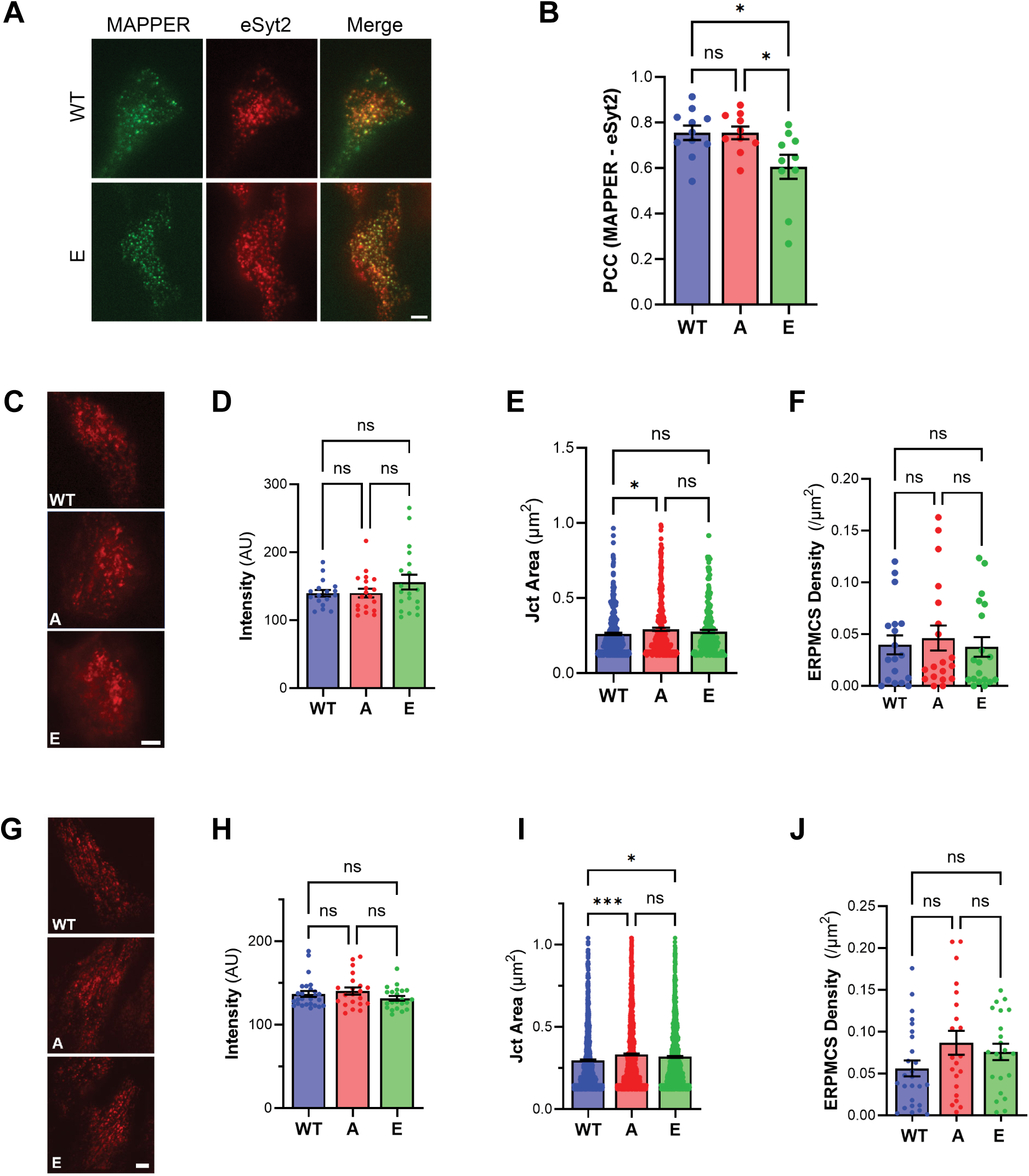
Functional characterization of the E-Syt2 mutants in interphase. (A) Representative TIRF images of E-Syt2 knockout HeLa cells transiently expressing GFP-MAPPER with either mCherry-E-Syt2-WT, or mCherry-E-Syt2-E mutant. Scale bar 5 µm. (B) Pearson’s correlation coefficient (PCC) analysis of the colocalization between MAPPER with either E-Syt2-WT, E-Syt2-A or E-Syt2-E mutants. Mean ± SEM; n = 10-11; Anova with Tukey correction; * p < 0.05, ns not significant. (C) Representative TIRF images of E-Syt2 knockout HeLa cells transiently expressing WT, E-Syt2-A or E-Syt2-E mutants using lipofectamine transfection. Scale bar 5 µm. (D-F) Quantification of ERPMCS. (D) Intensity per cell, n = 17-19. (E) Individual junction area, n = 281-372. (F) ERPMCS density per cell, n = 17-19. Mean ± SEM; Anova with Tukey correction; * p < 0.05, ns not significant. (G) Representative TIRF images of E-Syt2 knockout HeLa cells expressing WT, E-Syt2-A or E-Syt2-E mutant using adenoviral infection. Scale bar 5 µm. (H-J) Quantification of ERPMCS. (H) Intensity per cell, n = 21-26. (I) Individual junction area, n = 1076-1437. (J) ERPMCS density per cell, n = 21-26. Mean ± SEM; Anova with Tukey correction; * p < 0.05, *** p < 0.001, ns not significant.

We then transiently transfected the three constructs individually in the E-Syt2-KO cells (Fig. 2C) while controlling for expression levels at the cell level (Fig. 2D), and observed a modest increase in junction area with the E-Syt2-A but not the E-Syt2-E mutant (Fig. 2E). However, this effect did not translate when we quantified ERPMCS density, which was similar among the three clones (Fig. 2F).

We were concerned with the variability inherent in lipofectamine based transient transfection and thus generated adenoviruses to better standardize the expression of the three constructs (Fig. 2G and 2H). In this case we replicate the increase in junction area with the E-Syt2-A mutant (Fig. 2I) without any effect on the ERPMCS density (Fig. 2J). Increased junction area with the E-Syt2-A mutant would be consistent with increased tethering capacity but the observed changes despite being statistically significant were small.

Collectively these results suggest that the mitosis-specific phosphorylation of E-Syt2 has modest to no effects on its tethering capacity. Alternatively, given the redundancy in the E-Syts and other ERPMCS tethers, it is possible that the role of E-Syt2 phosphorylation was masked in interphase given the contribution of the other ERPMCS tethers, which ultimately determines ERPMCS area and density.

### E-Syt2 phosphorylation in mitosis

We had previously shown that the density of ERPMCS in mitosis decreases dramatically ^25^. This argues that the tethering capacity of all the ERPMCS tethers combined is reduced during mitosis. Thus, mitosis would provide a better context to assess the effect of phosphorylation on E-Syt2 tethering capacity as it can be quantified in the absence of the background tethering from the other ERPMCS tethers.

We infected E-Syt2-KO cells with the three E-Syt2 constructs and enriched for mitotic cells using nocodazole. It was difficult to quantify ERPMCS in the equatorial focal plane given their dearth (Fig. 3A), so we quantified them in the bottom focal plane of mitotic cells (Fig. 3B). The junction area was similar among the three constructs (Fig. 3C), but in this case we observed a robust rescue of the loss of ERPMCS in mitosis only when the E-Syt2-A mutant was expressed (Fig. 3D and 3E). This was apparent in the average density (Fig. 3D) and the density distribution histogram where the E-Syt2-A mutant had a marked shift to the right and a lower percent of cells with no junctions as compared to the E-Syt2-WT and E-Syt2-E expressing cells (Fig. 3E). This shows that mitotic cells expressing the non-phosphorylatable eSyt2-A mutant maintain more of their ERPMCS in mitosis compared to cells that express the wild type E-Syt2 or the E-Syt2-E mutant. This suggests that an E-Syt2 that cannot be phosphorylated in mitosis maintains its tethering potential.

**Figure 3.**
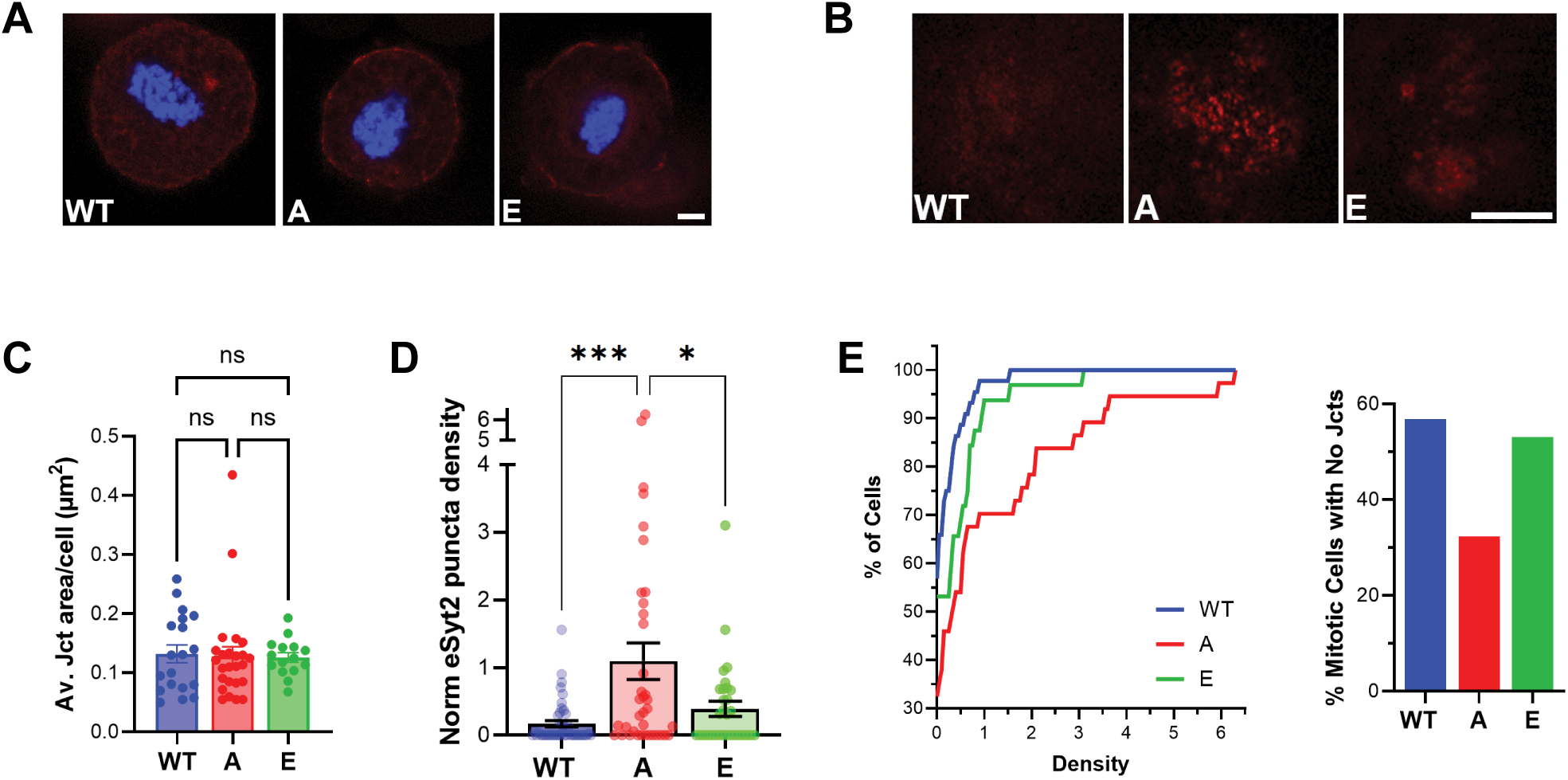
Non-phosphorylatable E-Syt2-A rescues ERPMCS loss in mitosis. (A-B) Representative confocal images of mitotic E-Syt2-KO cells expressing WT, E-Syt2-A, or E-Syt2-E at the equatorial (A) or bottom plane (B) with Hoechst DNA staining. Scale bar 5 µm. (C-D) Quantification of ERPMCS. (C) Average junction area per cell for the cells with junctions, n = 15-25. (D) Density of E-Syt2 positive junctions per cell, n = 32-44. Mean ± SEM; Anova with Tukey correction; * p < 0.05, *** p < 0.001, ns not significant. (E) Cumulative histogram distribution of the density of E-Syt2 positive ERPMCS in the population following expression of the three E-Syt2 constructs. The bar graph (right) shows the percentage of mitotic cells lacking detectable ERPMS.

A technical concern with the above results is that high levels of expression of tethers are known to enhance junction formation. In addition, although confocal imaging provides a good approximation of ERPMCS density, it is difficult to ascertain conclusively compared to the level of resolution obtained using electron microscopy (EM). To better control for these variables, we generated stable cells lines on the E-Syt2-KO background expressing either the wild-type protein or the E-Syt2-A mutant (Fig. S1C). We enriched for clones that express low levels of E-Syt2 by flow cytometry to avoid artificial junctions due to overexpression. We further treated the cells with nocodazole to enrich for mitotic cells and performed EM to quantify ERPMCS in mitosis. To avoid any bias in the quantification the entire dataset of 350 images was blinded and analyzed by four different individuals. Consistently we observed more ERPMCS in mitotic E-Syt2-A expressing cells compared to WT cells (Fig. 4). Example EM images of ERPMCS in mitosis are shown in Figure 4A for both the stable WT expressing cells and those expressing E-Syt2-A. Most of the detected junctions in mitosis had a relatively small junction length (Fig. 4A and 4B) with some instances in the cells expressing the E-Syt2-A mutant exhibiting a distended ER below the junction (Fig. 4A, double arrow). This may be indicative of the higher tethering capacity of the E-Syt2-A mutant in mitosis ‘pulling’ the ER to the PM.

**Figure 4.**
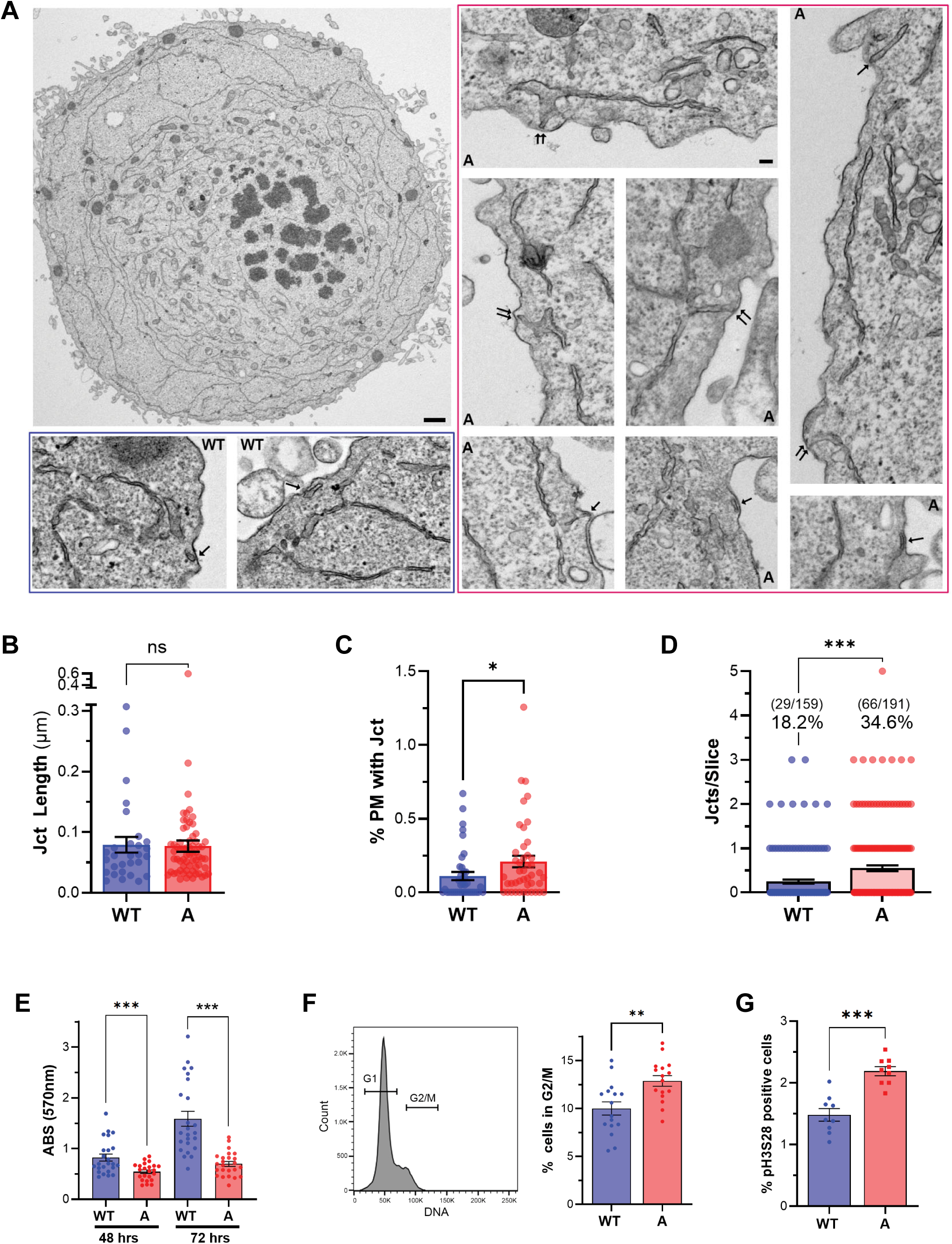
Electron microscopy quantification of ERPMCS in mitosis and effect on mitosis progression. (A) Representative section across a cell in mitosis at low magnification, with zoomed in examples of ERPMCS indicated by the arrows from stable cells expressing either E-Syt2-WT (WT) or E-Syt2-A mutant (A). Double arrows indicate junctions connected to a apparently distended ER. Scale bar: whole-cell image (E-Syt2-WT),1 µm; zoomed-in regions, 100 nm. (B–D) Quantification of ERPMCS. (B) junction length, n = 29-66. (C) percentage of the PM occupied by ERPMCS, n = 39-44. (D) Number of junctions per EM image, n = 159-191. *** p < 0.001, * p < 0.05, ns not significant. (E) MTT proliferation assay of E-Syt2 and E-Syt2-A stable cell line at 48 and 72 hours. Mean ± SEM; n = 23-24, *** p<0.001, unpaired t-test at each time point. (F) Flow cytometric cell cycle analysis of E-Syt2 and E-Syt2-A stable cells. (B) Example of the population cell cycle distribution following Hoechst nuclear staining(left). Quantification of the percentage of cells in the G2/M phase of the cell cycle (right). Mean ± SEM, n = 16, ** p<0.01, unpaired t-test. (G) Quantification of the percent of cells in mitosis in the stable E-Syt2 and E-Syt2-A cell lines. Mitotic cells are defined as those that stain positive for phospho-Ser28 histone H3. Mean ± SEM, n=9, *** p<0.001, unpaired t-test.

Mitotic cells stably expressing the E-Syt2-A mutant had significantly more ERPMCS as compared to those expressing E-Syt2-WT. The percent of the PM occupied by ERPMCS increased from 0.11±0.027 (n=39) in the WT cells to 0.21±0.04 (n=40) in the E-Syt2-A cells (Fig. 4C). The increase in ERPMCS was more pronounced when we quantified the number of contact sites per EM slice, where we again observed an increase in the percent of sections showing junctions in the E-Syt2-A images (34.6%; 66/191) as compared to WT (18.2%; 29/1159) (Fig. 4D). The average junctions per image detected were 0.25±0.046 in WT as compared to 0.55±0.065 in E-Syt2-A (Fig. 4D). Together the EM data support the conclusion from the confocal analyses, that expression of the E-Syt2-A non-phosphorylatable mutant in mitosis results in the persistence of ERPMCS that would otherwise be lost.

### E-Syt2-A cells exhibit defects in mitosis progression

The increase in ERPMCS we observed in the E-Syt2-A versus the E-Syt2-WT stable cells was consistent and significant, but rather modest. Although this aligns with the selection of stable cell lines that low levels of the E-Syt2 WT and A mutant, it made us question the physiological significance of such a moderate change. We therefore assessed cell growth and mitotic progression (Fig. 4E-G). Cell growth estimation using both the MTT (Fig. 4E) and Alamar blue assays (Fig. S1D) revealed a reduction in the E-Syt2-A cells compared to the WT cells. We further assessed the cell cycle distribution of both cell populations, which showed an increase in the percent of cells in the G2/M phase of the cell cycle in the E-Syt2-A cells (Fig. 4F). This suggested a delay in mitotic progression, which was confirmed by assessing the percent of cells in mitosis by staining for phosphorylation of histone H3 at Ser 28, a marker of mitotic cells (Fig. 4G). These data show that cells expressing E-Syt2-A progress through mitosis slower than the WT control cells.

## Discussion

Proper remodeling of organelles during mitosis has long been recognized as essential for division and the ensuing growth and differentiation. An interesting new area of investigation is how, concomitantly with this extensive remodeling, the connectivity and interaction between different organelles are altered in mitosis. Organelles maintain close connectivity through membrane contact sites ^11^; however, little is known about how these contact sites remodel in mitosis ^35^. Given the dramatic reorganization of most organelles in mitosis, one would expect that contact sites would also be affected. There is evidence for the remodeling of ERPMCS and MERCS in mitosis interestingly in opposite directions ^35^. MERCS expand in length in mitosis, which mediates better Ca^2+^ coupling between the two organelles presumably to support the increased energy demands of the dividing cell ^9,10^. MERCS expansion in mitosis is mediated by the moonlighting function of the Lamin B receptor (LBR) an inner envelope protein that supports nucleus structure in interphase but then redistributes to the ER upon nuclear envelope breakdown in mitosis and is enriched at MERCS ^10^. In contrast, ERPMCS are greatly reduced in mitosis, a loss that is associated with the significant reduction in Ca^2+^ influx through SOCE ^25^.

The molecular mechanisms controlling the loss of ERPMCS in mitosis are not known. We reasoned that phosphorylation during mitosis could play a role, given the well-defined central role of kinase cascades, particularly through CDK1 and MAPK, in driving cell division. Indeed, several of the known ERPMCS tethers are phosphorylated in a mitosis specific fashion (Table 1) ^32^. To test the role of phosphorylation in regulating tethering potential, we generated phosphomimetic and non-phosphorylatable mutants of E-Syt2 at the mitosis-specific sites. Careful quantification of ERPMCS localization of these mutants and their ability to modulate ERPMCS in interphase showed little differences (Fig. 1 and 2). But there were hints that phosphorylation may be playing a role. First, the phosphomimetic E-Syt2-E mutant had a lower colocalization index with the ERPMCS marker MAPPER (Fig. 2A-B); and second the area of ERPMCS was larger is cells expressing only the non-phosphorylatable E-Syt2-A mutant (Fig. 2E and 2I). We inferred that these small differences are likely due to the presence of the multitude of other tethers in interphase. So, we carried out the studies in mitosis as the tethering background from other tethers would be lower because of the inherent loss of ERPMCS. We did not observe any difference between the WT and E-Syt2-E in mitotic cells, but there was a pronounced increase in ERPMCS density and in the percent of mitotic cells with ERPMCS in cells expressing the syt2-A mutant (Fig. 3). These findings were confirmed by EM quantification of stable cells expressing either the WT or E-Syt2-A mutant at low levels to avoid overexpression artifacts (Fig. 4). Collectively these data show that inhibiting phosphorylation of E-Syt2 in mitosis preserves a higher tethering capacity. Interestingly, phosphorylation has been shown to modulate the function of another ERPMCS tether, TMEM24. In this case Ca^2+^-dependent phosphorylation releases TMEM24 from the PM in both insulin secreting cells and neurons ^36,37^. This suggests that tethers’ phosphorylation is a conserved mechanism to modulate their tethering capacity.

An important question is whether the loss of ERPMCS in mitosis is functionally significant. We had shown that the decrease in ERPMCS in mitosis is associated with the loss of SOCE and that forcing a constitutively active SOCE in mitosis leads to mitotic catastrophe and cell death ^25^. However, it is not clear whether the persistence of ERPMCS in mitosis is functionally consequential. The E-Syt2-A stable cell line offered a tool to test this as it had almost double the number of ERPMCS in mitosis compared to E-Syt2-WT and E-Syt2-E cells (Fig. 4). We therefore tested cell growth and cell cycle distribution and observed slower growth and an increased percent of the population in G2/M and specifically in mitosis in the E-Syt2-A cells (Fig. 4), arguing that E-Syt2-A cells progress through mitosis slower. This suggests that the loss of ERPMCS in mitosis is important to support timely cell division, where even the relatively small increase in the number of ERPMCS in mitosis in the E-Syt2-A mutant was associated with slower cell division.

Cobb et al. had shown that when the neuronal potassium channel Kv2.1 is expressed in COS-1 cells it localizes to ERPMCS in a mitosis-specific fashion ^38^. This would argue that mitotic COS-1 cells maintain ERPMCS in mitosis. Our results in HeLa cells show a dramatic reduction of ERPMCS in mitosis ^25^. The reasons for these differences are not clear but could be due to differential regulation in distinct cell types, the overexpression of Kv2.1 which could stabilize junctions, or the nature of the tethers tested.

In summary, our results support a model where mitosis specific phosphorylation of the ERPMCS tether E-Syt2 reduces its tethering capacity and thus contributes to the loss of ERPMCS observed in mitosis. Interesting, partial reversion of this loss is associated with delayed progression through mitosis. This suggests that phosphorylation of ERPMCS tethers in mitosis is an important regulatory mechanism controlling ER and PM association.

## Methods

### Plasmids

GFP-MAPPER, mCherry-E-Syt2 (E-Syt2-WT), TMEM24-mCherry, mCherry-ORP8, GFP-VAPB, and mCherry-GRAMD1A were generous gifts from the Liou ^34^, De Camilli ^27,37^, Balla ^39^, Trimmer ^40^, and Nunnari ^41^ Lab, respectively. Non-phosphorylatable mutant plasmid E-Syt2-A mutant (S612A, S623A, T629A, S667A, T677A) and the phosphomimetic plasmid E-Syt2-E mutant (S612E, S623E, T629E, S667E, T677E) were synthetized and sequence confirmed by Gene Universal. Adenoviral particles expressing mCherry-E-Syt2-WT, -A and -E mutants were produced by Vector Biolabs. HeLa cells were infected with recombinant adenovirus at a multiplicity of infection (MOI) of 100 PFU/cell. The plasmids pLenti-CMV-GFP-2A-Puro-mCherry-E-Syt2-WT and pLenti-CMV-GFP-2A-Puro-mCherry-E-Syt2-A were generated by subcloning E-Syt2-WT and E-Syt2-A mutant into pLenti-CMV-GFP-2A-Puro vector (GenScript).

### Cell Lines and Treatment

HeLa and HEK293T cells (ATCC) were cultured in Dulbecco’s Modified Eagle’s Medium (DMEM) (Gibco) supplemented with 10% Fetal Bovine Serum (FBS) and 1% penicillin-streptomycin antibiotic mixture, maintained at 37°C in a humidified incubator with 5% CO₂.

E-Syt2-knockout (KO) HeLa lines were generated using pLentiCRISPR all in one lentiviral vector system (GenScript). Briefly, two sgRNAs targeting human *E-Syt2* (sgRNA1: 5′-TAGTTTCACGAAGGTCGACG-3′; sgRNA2: 5′-AGTGGAACGGTGATTCGATT-3′) were cloned into the pLentiCRISPR vector (GenScript). Lentiviral particles were produced in HEK293T cells by co-transfecting the transfer constructs with standard packaging plasmids, following the manufacturer’s instructions and used to transduce HeLa cells. Following puromycin selection, single-cell clones were isolated, expanded, and validated for E-Syt2 knockout by Western blot.

To establish the E-Syt2-WT and E-Syt2-A stable cell lines, lentiviral particles were produced in HEK293T cells by co-transfecting either pLenti-CMV-GFP-2A-Puro-mCherry-E-Syt2-WT or pLenti-CMV-GFP-2A-Puro-mCherry-E-Syt2-A with standard packaging plasmids, following the manufacturer’s instructions and used to transduce E-Syt2 KO HeLa cells. Single cell clones were isolated by single cell sorting into 96-well plate using BD FACS Aria III cell sorter. GFP positive cell clones were expanded and validated for the expression of mCherry-E-Syt2-WT and mCherry-E-Syt2-A by TIRF microscopy and Western blot.

For mitotic cell enrichment, the cells were treated with 400 ng/mL of Nocodazole (Sigma-Aldrich) for 18 hr. For transient transfection, cells were transfected with lipofectamine 2000 reagent (Life Technologies) following the manufacturer’s protocol.

### Western Blotting

Cells were lysed in ice-cold RIPA buffer supplemented with protease and phosphatase inhibitors. Lysates were incubated on ice for 30 min, centrifuged at 12,000 rpm for 15 min at 4 °C, and supernatants collected. Protein concentrations were determined by BCA assay. Samples were analyzed by SDS–PAGE followed by Western blotting using anti-E-Syt2/FAM62B polyclonal antibody (PA5-109710; Invitrogen), anti-E-Syt2 polyclonal antibody (HPA002132; Sigma) or anti-E-Syt2 polyclonal antibody (24385-1-AP; Proteintech).

### Microscopy

Cells were plated in 35 mm poly-D-lysine coated glass-bottom dishes (MatTek). Prior for imaging, cells were stained with Hoechst 33342 (2μg/mL; Life Technologies) for 20 minutes at 37°C and then fixed with 4% paraformaldehyde.

Total Internal Reflection Fluorescence (TIRF) images were acquired using a Zeiss Z1 Axio Observer inverted microscope with an alpha Plan-Apochromat 63×/1.46 Oil Corr M27 TIRF objective. An epifluorescence image was also taken alongside TIRF image.

Mitotic cells were imaged on a Zeiss LSM 880 confocal microscope using a Plan Apo 63×/1.4 oil DIC II objective with the pinhole set at 1 AU. For each mitotic cell two images were captured one at the bottom and the second at the cell equator.

Images were analyzed using Zeiss ZEN Black and Image J (FIJI) software. Quantification of ERPMCS was performed using a macro developed in FIJI to automate the process and minimize bias. The LUT was normalized based on the maximal expression for each experiments. Background subtraction was performed using a rolling ball of 5. Images were then automatically thresholded using the ImajeJ built in ‘Intermodes’ algorithm with the threshold set at the same levels for all images of the three genotypes for each experiment. The analysis was repeated on at least 3 biological replicates.

### Electron microscopy

Cell pellets were fixed overnight at 4°C in EM fixation solution containing 4% PFA and 2.5% glutaraldehyde (Electron Microscopy Sciences) in 0.1 M Na-cacodylate buffer (pH 7.4). Samples were then washed three times with 0.1 M Na-cacodylate buffer and post-fixed with 1% OsO4– 1.5% K-ferricyanide in 0.1 M Na-cacodylate buffer at room temperature for 1 h. Following three washes with buffer, samples were dehydrated through a graded ethanol series and embedded in EMbed 812 resin (Electron Microscopy Sciences Cat. #14900) according to the manufacturer’s instructions. Ultrathin sections (70 nm) were collected, contrasted with lead citrate, and examined using a JEM 1400 transmission electron microscope (JEOL USA, Inc.) operated at 100 kV. Digital images were acquired using a Veleta 2K × 2K CCD camera (Olympus-SIS).

### Proliferation assay and cell cycle analysis

Cells were seeded in 96-well plates at a density of 5,000 cells per well. Cell proliferation was assessed at 48 and 72 hours using either the MTT assay (Sigma-Aldrich) or the AlamarBlue assay (Invitrogen). For the MTT assay, cells were incubated with 5 mg/mL MTT reagent for 1 hour, after which the resulting formazan crystals were dissolved in dimethyl sulfoxide (DMSO) prior to absorbance measurement at 570 nm with a reference wavelength of 690 nm. For the AlamarBlue assay, the reagent was added directly to the culture medium at the recommended dilution and incubated for 4 hours without medium replacement. Absorbance was measured at 570 and 600 nm, and values at 570 nm were corrected by subtracting the background signal from growth medium alone. Data were acquired using a CLARIOstar Plus plate reader (BMG Labtech). Each experimental group contained seven technical replicates.

Cells were stained with Hoechst 33342 (Invitrogen, H3570) and Alexa Fluor 647-conjugated rat anti-histone H3 (pS28) antibody (BD Phosflow, 558217) for cell cycle analysis, following the manufacturer’s instructions. Flow cytometry data were acquired and analyzed using FlowJo v10 software (Tree Star).

### Statistics

Graphpad PRISM was used for statistical analyses and to generate graphs. Data are presented as Mean ± SEM with the statistical test and significance as indicated in the Figure legend.

## Data availability

All data are included in the manuscript and Supplementary Information, or available from the authors. Unique reagents generated during this work are available to colleagues in the scientific community upon request. Please contact the corresponding author at.

## Acknowledgments

This work is funded by grant # ARG 01-0517-230201 from the Qatar Research, Development and Innovation Council. The findings herein reflect the work and are solely the responsibility of the authors. Partial support for the work as well as the Weill Cornell Medicine Qatar Cores came from the Biomedical Research Program (BMRP), a program funded by Qatar Foundation. We are grateful to the Microscopy Core at WCMQ for its exceptional support for this study.

## Authors Contributions

J.A.H and F.Y. performed experiments and analyzed data, K.S.K and L.Z. performed experiments. K.M. conceptualized and supervised the project, led study design, analyzed data, secured funding and wrote the manuscript.

---

**Supplemental Figure 1.**
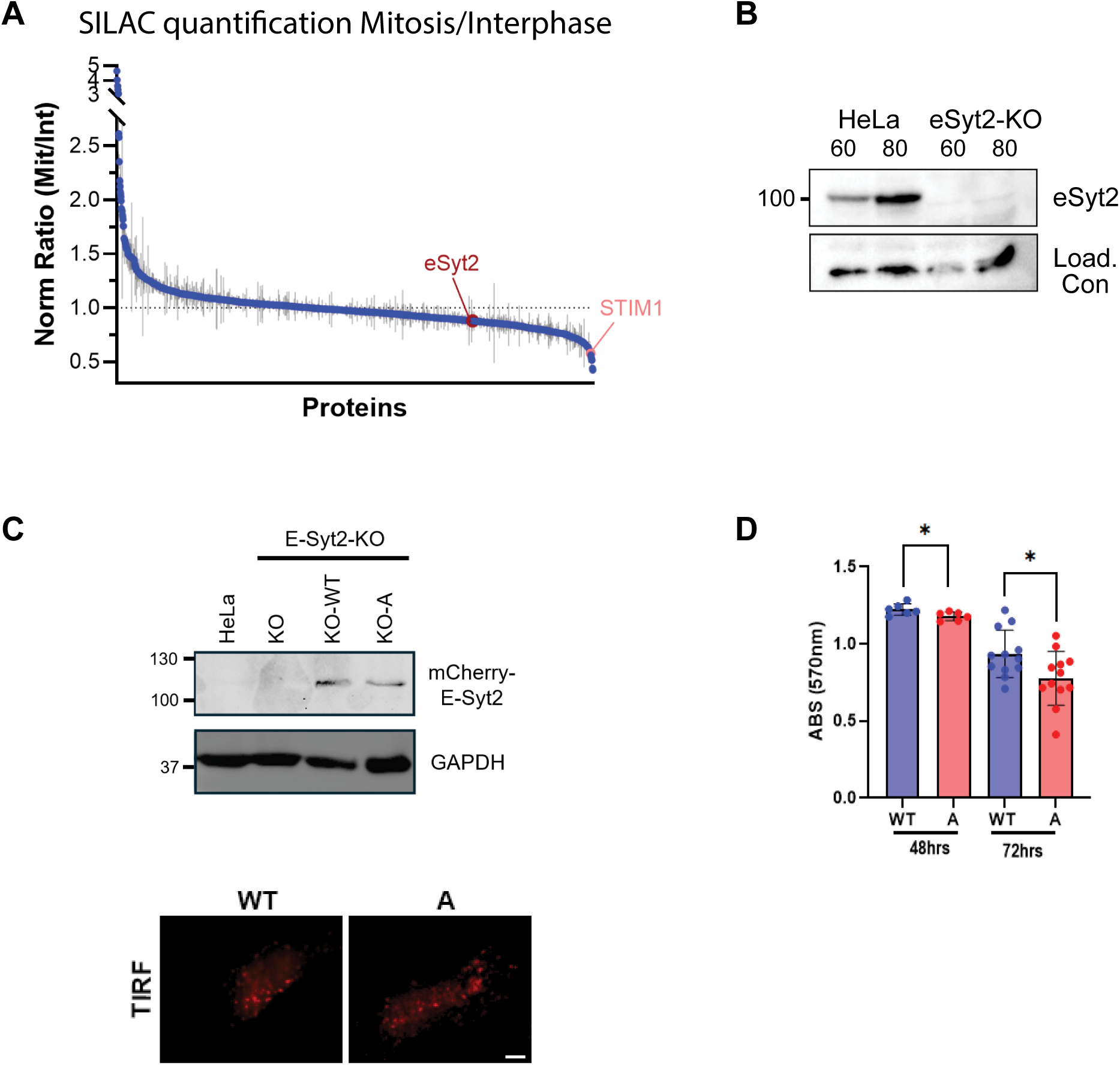
(A) SILAC-based quantification of protein abundance in mitosis as compared to interphase. (B) Western blot validation of E-Syt2 knockout HeLa cell line using the Sigma anit-E-Syt2 antibody (HPA002132). (C) WB validation of the E-Syt2-KO cell lines stably expressing the WT E-Syt2 or the E-Syt2-A mutant (top). TIRF images of the stable E-Syt2-KO cell lines expressing the WT protein or the E-Syt2-A mutant. (D) AlamarBlue proliferation assay of E-Syt2 and E-Syt2-A expressing cells at 48 and 72 hours. Mean ± SEM; n=6-12. * p < 0.05, unpaired t-test at each time point.

**Supplemental Figure 2.**
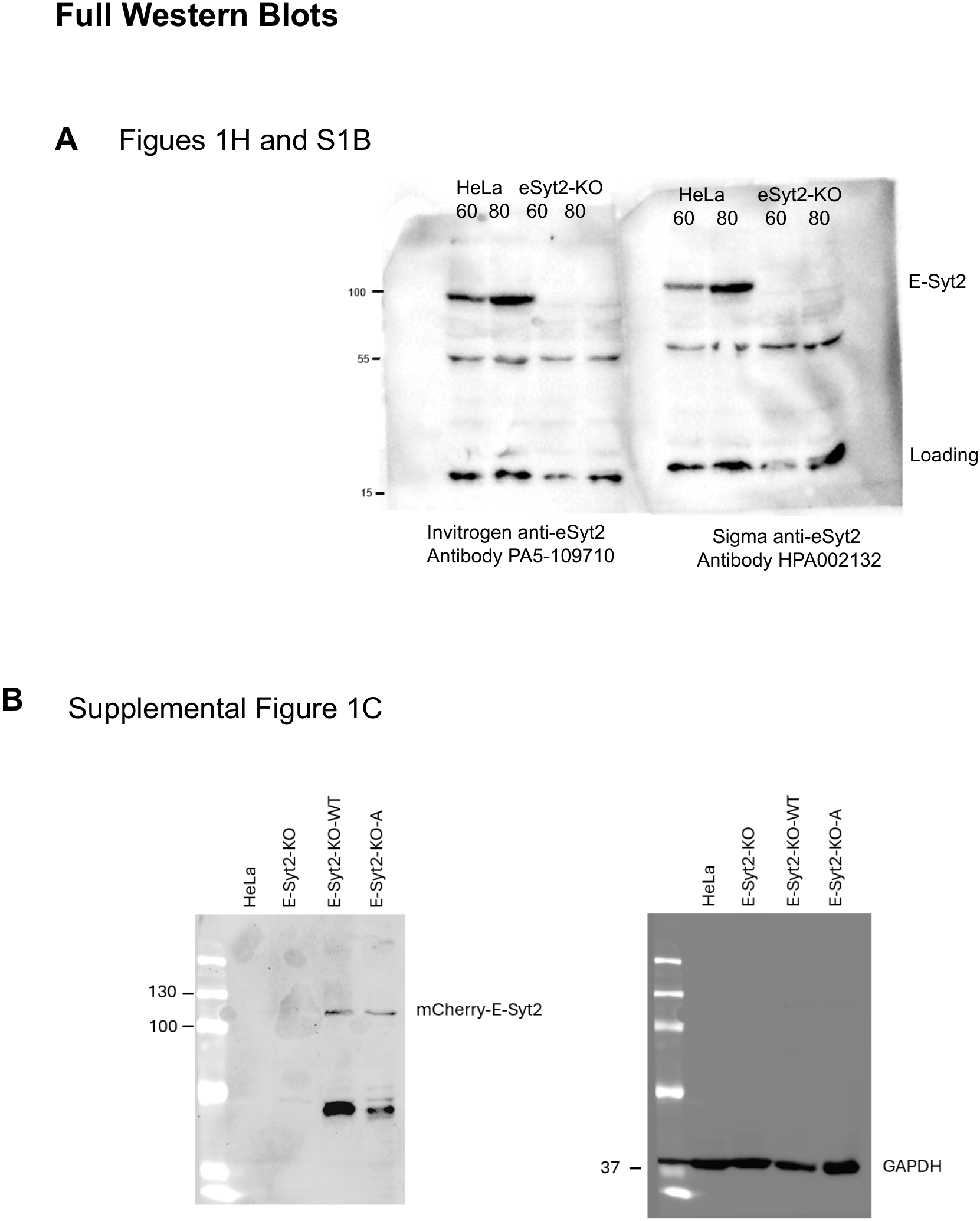
Original uncropped WB images as labelled.

